# Targeting long non-coding RNA RP11-502I4.3 delays the trend of angiogenesis in diabetic retinopathy

**DOI:** 10.1101/2024.10.15.618482

**Authors:** Lan Zeng, Yuhao Wu, Lijuan Zhu, Junhao He, Yuan Yuan, Xiaocong Wang, Kai Tang, Wei Tan

## Abstract

Based on our previous findings, we hypothesized that the long non-coding RNA RP11-502I4.3 may be involved in angiogenesis associated with Diabetic retinopathy (DR). We investigated the role of RP11-502I4.3 in DR by examining its regulation of vascular endothelial growth factor (VEGF). We assessed differences in RP11-502I4.3 expression between the normal control group and streptozotocin-induced diabetic rats or high glucose (HG)-stimulated human retinal microvascular endothelial cells (HRMECs). VEGF expression was measured with and without lentiviral vectors overexpressing RP11-502I4.3. We analyzed the structural and functional alterations related to DR. Our analysis revealed that RP11-502I4.3 expression was lower in the retinas of diabetic rats and in HG-stimulated HRMECs compared with normal glucose conditions. Overexpressing of RP11-502I4.3 resulted in decreased VEGF levels. Diabetic rats exhibited retinopathy characterized by thinning of the retinal layer thickness, structural changes in the inner and outer nuclear layers, a reduced count of retinal ganglion cells, and the presence of acellular capillaries. The proliferative activity, migration count, and tube formation ability of HG-treated HRMECs were significantly higher than those of the normal control group; however, these changes were delayed by RP11-502I4.3 overexpression. RP11-502I4.3 downregulation in DR appears to promote angiogenesis.

## Introduction

Diabetes mellitus is a significant public health problem[1]. Diabetic retinopathy (DR), a neurovascular disorder caused by chronic hyperglycemia, is a leading cause of blindness worldwide[2]. Proliferative diabetic retinopathy (PDR) is characterized by angiogenesis that begins in the retina and can extend into the vitreous, driven by increased vascular endothelial growth factor (VEGF)[3]. Retinal neovascularisation is prone to bleeding and fibrosis, which may result in vitreous hemorrhage and retinal detachment, ultimately leading to vision loss[4]. Treatment outcomes for PDR are often suboptimal[5]. Therefore, detailed studies on the pathogenesis of PDR are essential.

Long non-coding RNAs (lncRNAs) regulate gene expression and protein production[6]. Many lncRNAs have been associated with vascular oculopathies and proliferative retinopathy[7, 8]. With the advent of new RNA therapeutic technologies, treatments targeting lncRNAs may offer a promising new approach for PDR.

Recently, high-throughput sequencing has significantly accelerated the finding of novel lncRNAs and their biological functions. Yan *et al*. were the first to report that 303 lncRNAs were differentially expressed in the retina of early DR, with 89 upregulated and 214 downregulated[9]. Among these, MALAT1 expression was notably upregulated[9]. Other lncRNAs associated with DR, such as ANRIL[10], MIAT[11], MEG3[12], H19[13], PVT1[14], BANCR[15], and NEAT1[16] have also been reported. However, the function of lncRNA in PDR has yet to receive much attention.

We analyzed a microarray dataset (GSE191210) and found that lncRNA RP11-502I4.3 expression was significantly lower in the vitreous humor (VH) of patients with PDR compared with those with idiopathic macular hole (IMH)[17]. Based on these findings, we speculated that RP11-502I4.3 may be involved in angiogenesis in PDR. We investigated the expression levels and functions of RP11-502I4.3 in human retinal microvascular endothelial cells (HRMECs) exposed to high glucose (HG) and in the retinal tissues of streptozotocin (STZ)-induced diabetic rats.

## Materials and Methods

### Animals

All rats were handled following the Association for Research in Vision and Ophthalmology Statement for the Use of Animals in Ophthalmic and Vision Research. The animal study was approved by the Ethics Review Committee of First People’s Hospital of Zunyi (Zunyi, China; project number 2021-2-31). Adult male Sprague-Dawley (SD) rats were housed under specific pathogen-free conditions at the Animal Experimental Centre of Zunyi Medical University. The rats were divided into four experimental groups: a normal control group (right and left eyes of eight rats), a diabetes group (right and left eyes of eight rats), an overexpression control group (left eyes of 16 rats), and an overexpression group (right eyes of 16 rats).

To induce diabetes, the rats were fed a high-fat diet for four weeks. The rats received an intraperitoneal injection of STZ (Solarbio, China; 40 mg/kg in citrate buffer, pH 4.3) after fasting for 16 hours, whereas the normal control group received equivalent volumes of citrate buffer. After one week of STZ induction, diabetes was confirmed by blood glucose levels exceeding 16.7 mmol/L. In addition, RP11-502I4.3 overexpression models were established. All animals were monitored for changes in body weight and blood glucose levels. Twelve weeks after diabetes induction, tissues were collected from all rats.

### Intravitreal injection

The lentiviral vector (LV) was designed and synthesized by Hejin Company (Guizhou, China). Diabetic rats were anesthetized with an intramuscular injection of ketamine at a dose of 30 mg/kg. Tropicamide eye drops were then applied to dilate the pupils. Using a 33G microsyringe (Thermo, USA), we injected 1 µL of RP11-502I4.3 overexpression virus into the right vitreous cavity. Similarly, the control virus was injected into the left vitreous cavity of these rats. Finally, successful RP11-502I4.3 overexpression was confirmed using quantitative real-time polymerase chain reaction (qRT-PCR).

#### qRT-PCR

TRIzol reagent (Thermo Scientific, USA) was used to extract RNA from the retinas of rats and HRMECs. The RNA concentration was measured using a NanoDrop LITE spectrophotometer (Thermo Scientific, USA). Total RNA was reverse transcribed using the PrimeScrip™ RT Reagent Kit Reverse Transcription System (TaKaRa, Japan). Transcript levels were determined using the 2× SYBR Green qRT-PCR Mix (Solarbio, China) and a CFX96 real-time quantitative PCR detection system (Bio-Rad, USA). Relative gene expression was calculated using the 2^-ΔΔCt^ method, with β-actin serving as the internal control. The primer pairs used are listed in Table 1.

### Haematoxylin and eosin (H&E) staining

The eyeballs of the autopsied rats were placed into clean 1.5 mL Eppendorf tubes and fixed in 4% paraformaldehyde (Solarbio, China) at 37°C for more than 24 hours. The corneas, lenses, vitreous, and extraocular muscles were then removed. After washing in phosphate-buffered saline (PBS; Solarbio, China) and dehydrating in an alcohol gradient, the eyeballs were cleared in xylene I, II, and III, followed by immersion in paraffin, embedding, sectioning (5 µm), and baking. The paraffin sections were deparaffinized in xylene I and II, rinsed in anhydrous alcohol, and placed in an alcohol gradient. Follow-up steps follow the conventional H&E staining procedure (Solarbio, China). Retinal thickness and the number of retinal ganglion cells (RGCs) were measured at 0.8 mm from the optic nerve.

### Periodic acid-Schiff (PAS) staining

At autopsy, each eyeball was fixed as previously described. The retinas were isolated under a dissecting microscope (Olympus, Japan) and thoroughly washed. The retinas were incubated in a 3% trypsin solution (Solarbio, China). Isolated retinal vessels were stained with PAS (Solarbio, China) and visualized.

### Fluorescence in situ hybridization (FISH)

Tissue fixation, dehydration, sectioning, and deparaffinization were performed as previously described. Antigen retrieval was conducted in 0.01 mmol/L citric acid buffer at 100 °C for 15 minutes. After cooling, the sections were rinsed in PBS three times and incubated with the pre-hybridization solution (RiboBio, China) in a moisture box at 37°C for one hour. Subsequently, the pre-hybridization solution was removed, and the sections were incubated with the hybridization solution (RiboBio, China) at 37°C overnight. The hybridization solution was then rinsed off. Following counterstaining with DAPI (Beyotime, China), images were obtained using a fluorescence microscope (Olympus, Japan).

### Cells culture

In this study, HRMECs were purchased from the Beijing Beina Chuanglian Biotechnology Research Institute. The cells were divided into four groups: a normal control group (no intervention), an HG group (30 mmol/L glucose added to the medium and cultured for 48 hours), an overexpression control group, and an overexpression group. HRMECs were cultivated in a complete endothelial cell medium (ECM; ScienCell, USA) in an incubator (5% CO_2_ at 37°C).

### Cells transfection

The sequence used to overexpress RP11-502I4.3 was designed and synthesized by Hejin Company (Guizhou, China). The cells were inoculated into new six-well plates and cultured in an incubator for approximately 24 hours. Transfections were performed according to the instructions of the manufacturer (Hejin, China). The transfection efficiency reached 70-75% after 72 hours, as assessed using fluorescence microscopy (Olympus, Japan). The experiment was repeated three times in parallel.

### Cell Counting Kit-8 (CCK-8) assay

The cells were seeded into a 96-well plate at a density of 2 × 10^5^ cells/mL. After 48 hours of incubation, CCK-8 reagent (Beyotime, China) was added to each well at a ratio of 10:1. The plates were then incubated for one hour. Cell viability was assessed by measuring the absorbance of the cells at 450 nm using an i3*x* multifunctional enzyme labeling detector (Molecular Devices, USA). The experiment was repeated three times in parallel.

### Transwell assay

HRMECs were digested with trypsin, diluted to a density of 3 × 10^5^ cells/mL in complete ECM, and inoculated into the chambers of a 24-well Transwell plate. 500 µL of complete ECM was added to the lower chambers, and the cells were cultured for 48 hours. After this incubation, the lower chambers were fixed for 20 minutes. A 0.1% crystal violet solution (Beyotime, China) was then added to the chambers for 25 minutes for staining. The stained cells were counted using an upright microscope (Olympus, Japan). The experiment was repeated three times in parallel.

### Tube formation assay

A 50 µL/well Matrigel matrix (Corning, USA) was added to a 96-well plate, taking care to avoid bubble formation. The plate was then placed in a cell incubator (5% CO_2_ at 37°C) for one hour to allow for solidification. Cells from the four groups were then inoculated at a density of 3 × 10^5^ cells/mL into the 96-well plate. After incubation, the plate was removed and observed under a microscope (Olympus, Japan). The experiment was repeated three times in parallel.

### Statistical analysis

The above experiments were repeated three times in parallel. We used Image J 1.8.0 software for data analysis and GraphPad Prism 8 software for statistical analysis. Comparisons between two groups were performed using independent-samples *t*-tests. For comparisons involving more than two groups, a one-way analysis of variance was employed. When the data were not normally distributed, the rank-sum test was used. Statistical significance was set at *p* < 0.05.

## Results

### Differences in RP11-502I4.3 expression in a diabetic rat model

In a previous study [17], we performed microarray-based analysis to compare the lncRNA expression profiles between patients with PDR and those with IMH (Fig. 1A). The expression levels of RP11-502I4.3 in the VH of patients with PDR were significantly lower, as confirmed by qRT-PCR (Fig. 1B).

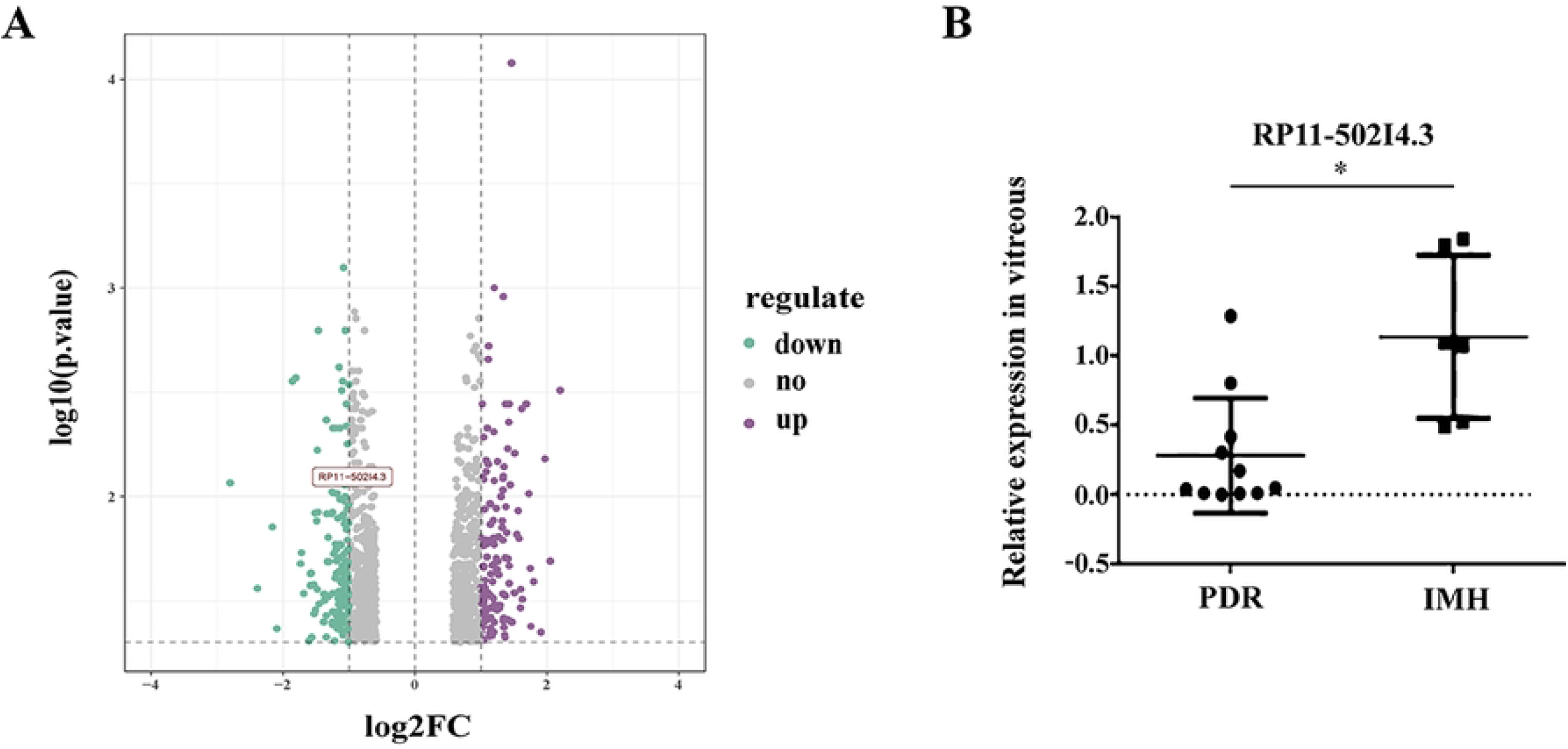

However, the involvement of RP11-502I4.3 in PDR remains unclear. We established a diabetic rat model to verify its expression. After being fed a high-fat diet for four weeks, adult SD rats were injected intraperitoneally with STZ (Fig. 2A). We measured rats’ blood glucose levels and body weights 12 weeks after STZ injection. The blood glucose levels in the diabetes group were significantly higher than those in the normal control group, and body weight was notably reduced (Fig. 2B and C). Indicators of retinopathy in diabetic rats included reduced retinal layer thickness (RLT) and retinal cell count[18]. In this study, we observed thinning of the RLT, structural changes in the inner nuclear layer (INL) and outer nuclear layer (ONL), and a decreased count of RGCs in diabetic rats compared with non-diabetic rats (Fig. 2D-F). Acellular capillaries (ACs) were also present in the retinal tissue of diabetic rats (Fig. 2G and H).

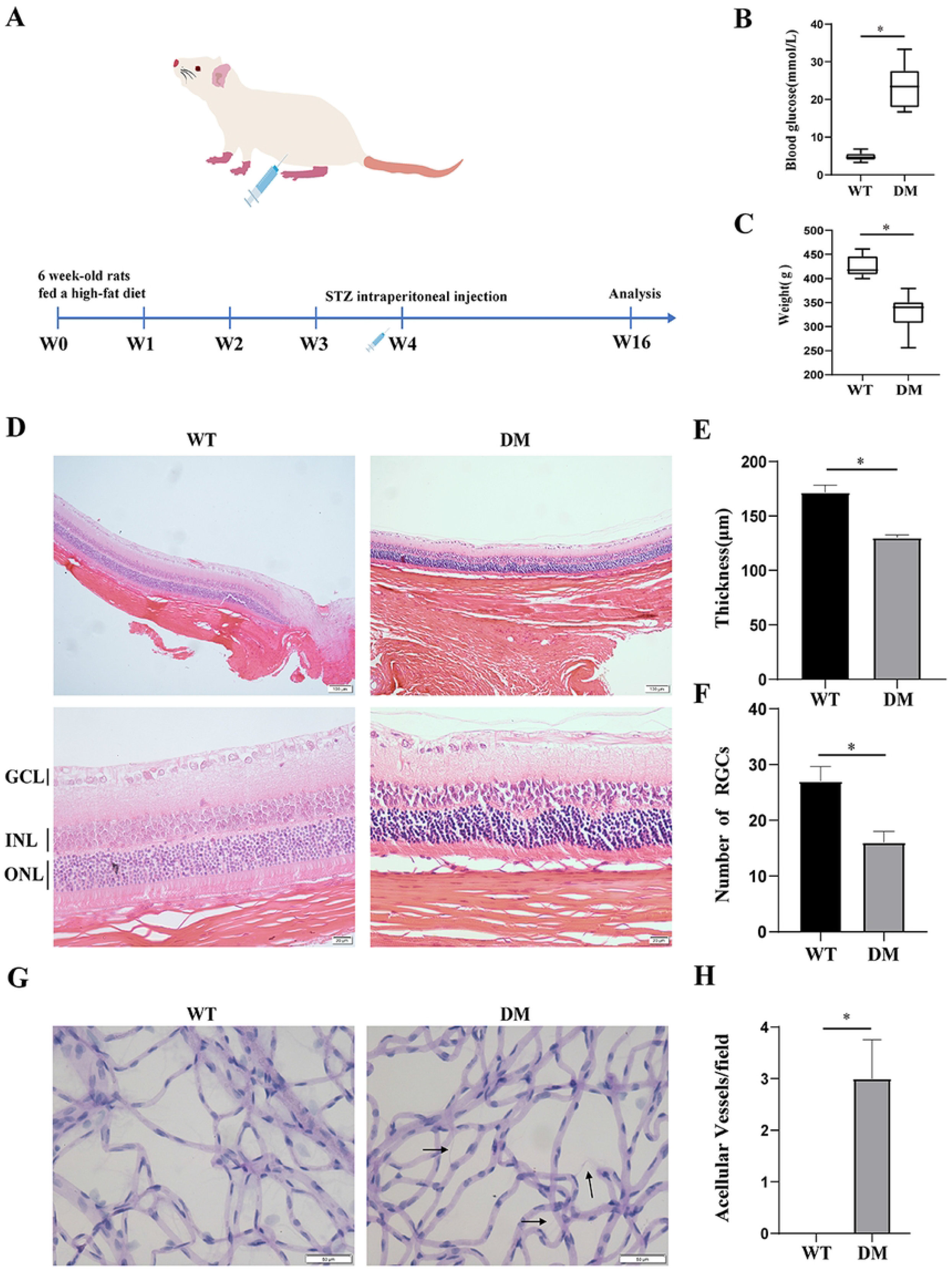

VEGF plays a vital role in the development of DR, and its increase drivers of retinal neovascularization[19]. Accordingly, we measured VEGF gene expression and found that it was significantly increased in the retinal tissue of diabetic rats (Fig. 3A). In addition, RP11-502I4.3 expression was decreased in the retinas of diabetic rats compared with non-diabetic rats (Fig. 3B), and it was predominantly localized in the cytoplasm (Fig. 3C).

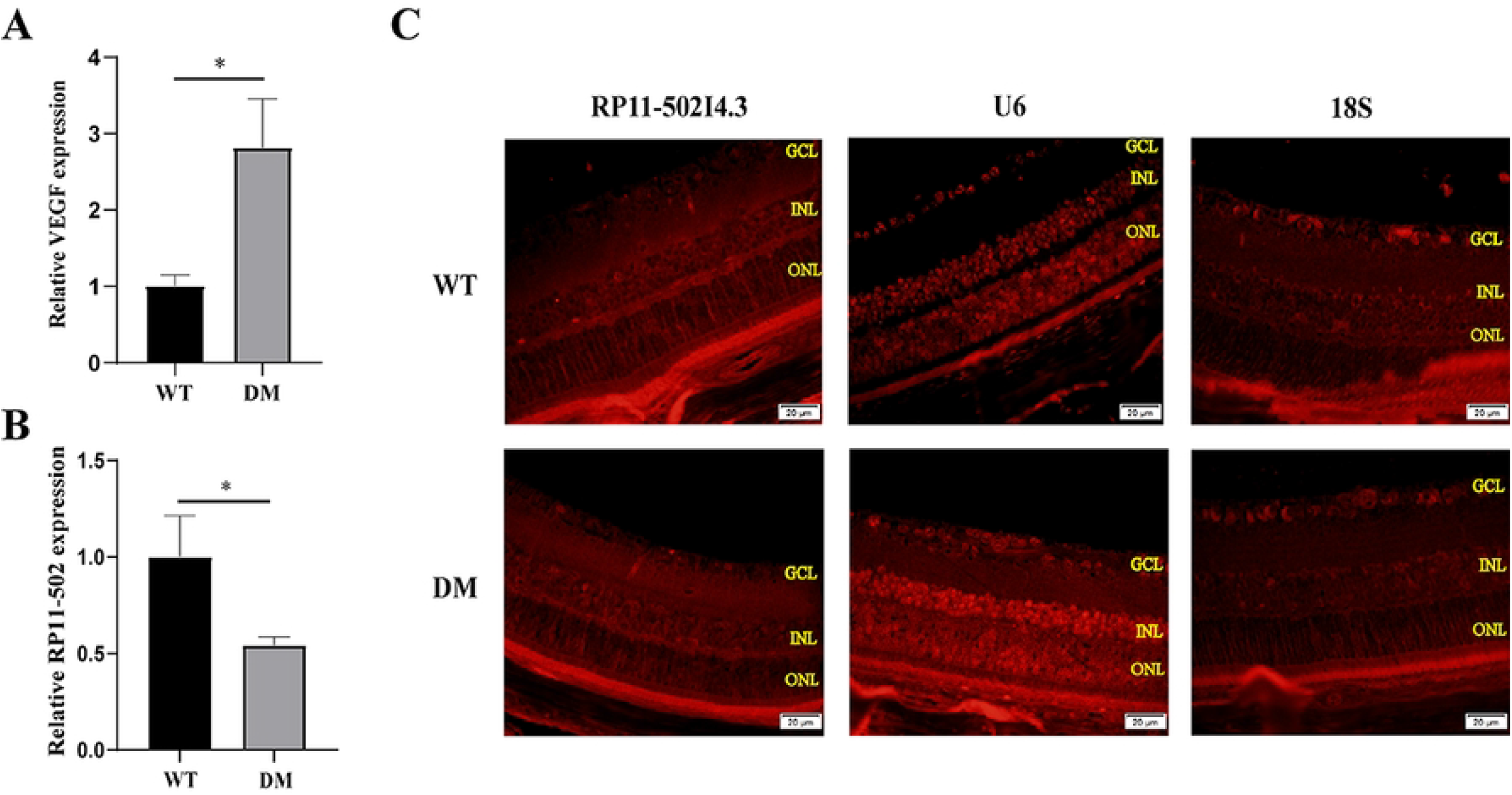

### Establishment of RP11-502I4.3 overexpression models

*In vivo* experiments, an overexpression model of RP11-502I4.3 was established using a LV to investigate the relevance of RP11-502I4.3 downregulation in neovascularisation (Fig. 4A and B). Diabetic rats were selected for intravitreal injection one week after STZ induction (Fig. 4C). Verification was conducted using qRT-PCR and FISH (Fig. 4D and E). The expression levels of RP11-502I4.3 in the overexpression group were significantly higher than those in the overexpression control group (Fig. 4D).

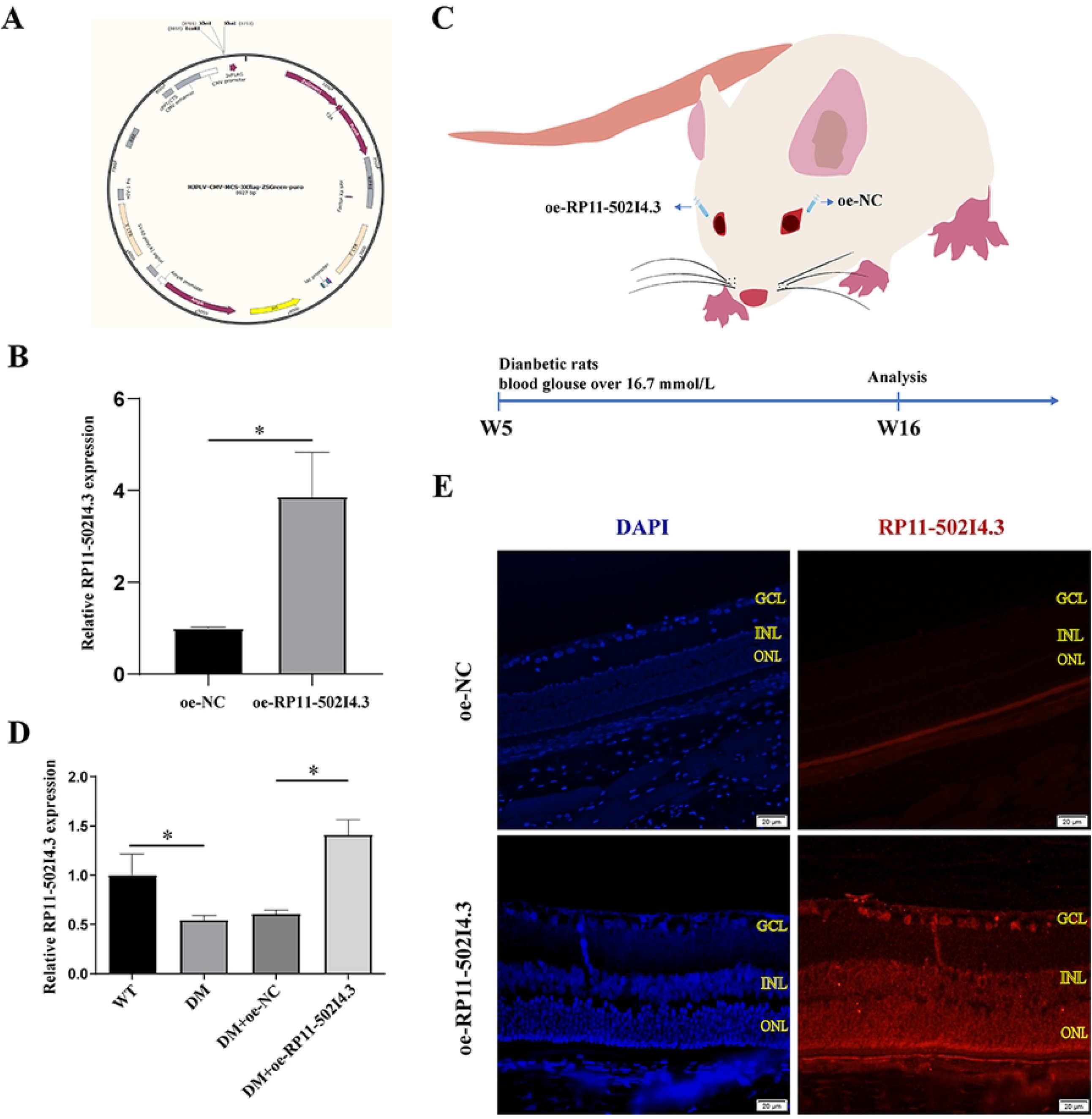

Next, we developed a cell model for RP11-502I4.3 overexpression (Fig. 5A and B). The transfection efficiency of overexpression was evaluated 72 hours after lentivirus-mediated transfection. Both the overexpression and negative control groups were assessed for the expression of green fluorescent protein, with transfection efficiency exceeding 70% (Fig. 5C). The verification process was performed similarly (Fig. 5D).

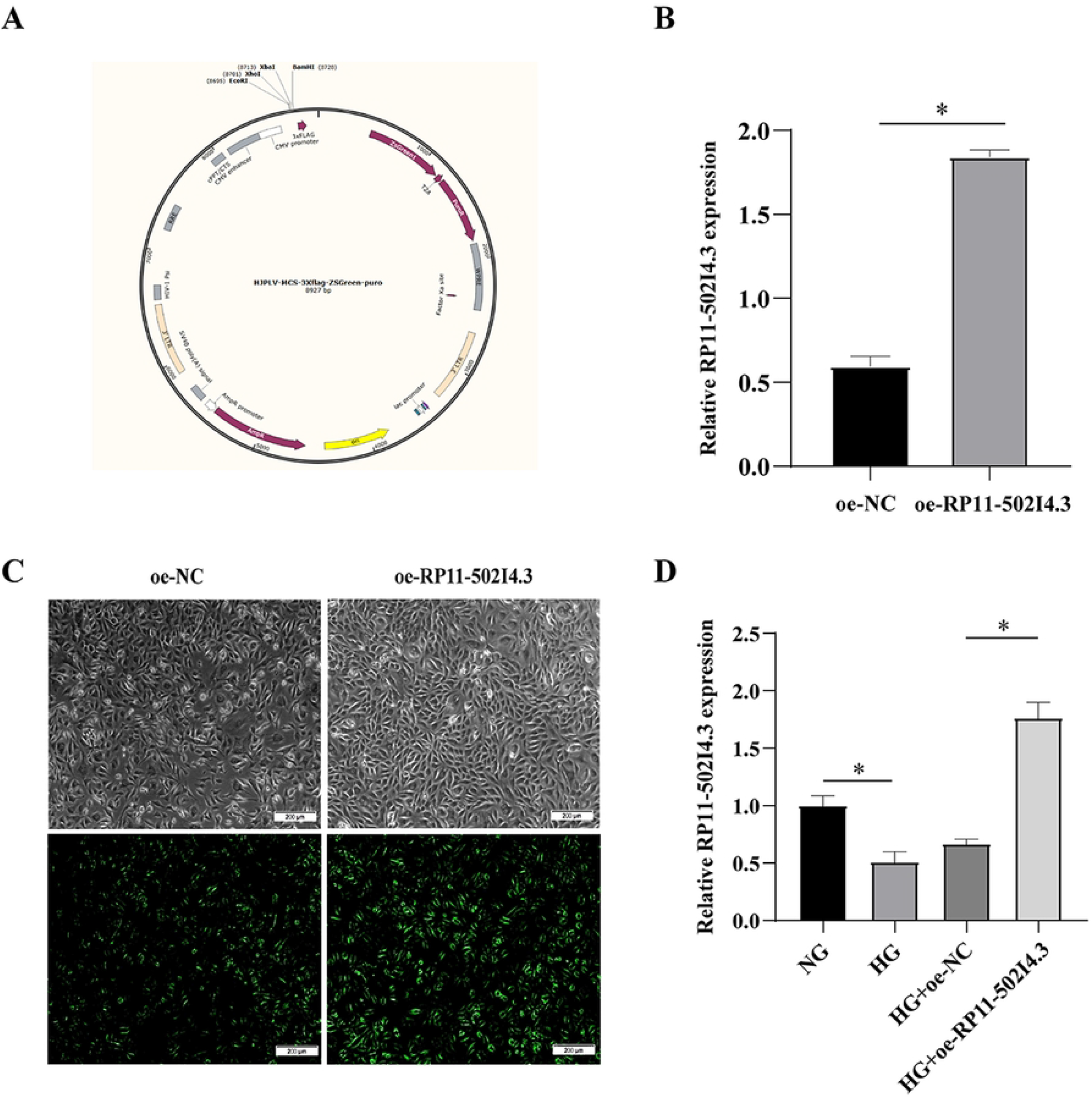

### RP11-502I4.3 Overexpression delayed the trend of retinopathy in the diabetic rat model and angiogenesis HG-treated HRMECs model

Using H&E staining, we observed the occurrence of retinopathy in diabetic rats, including thinning of the RLT, structural changes in the INL and ONL, and a decreased count of RGCs. In contrast, diabetic rats overexpressing RP11-502I4.3 showed delayed progression of these changes (Fig. 6A-C). The number of ACs in diabetic rats with RP11-502I4.3 overexpression was significantly lower than that in diabetic rats without RP11-502I4.3 overexpression (Fig. 6D and E). In addition, VEGF gene expression was significantly downregulated in diabetic rats after RP11-502I4.3 overexpression (Fig. 6F).

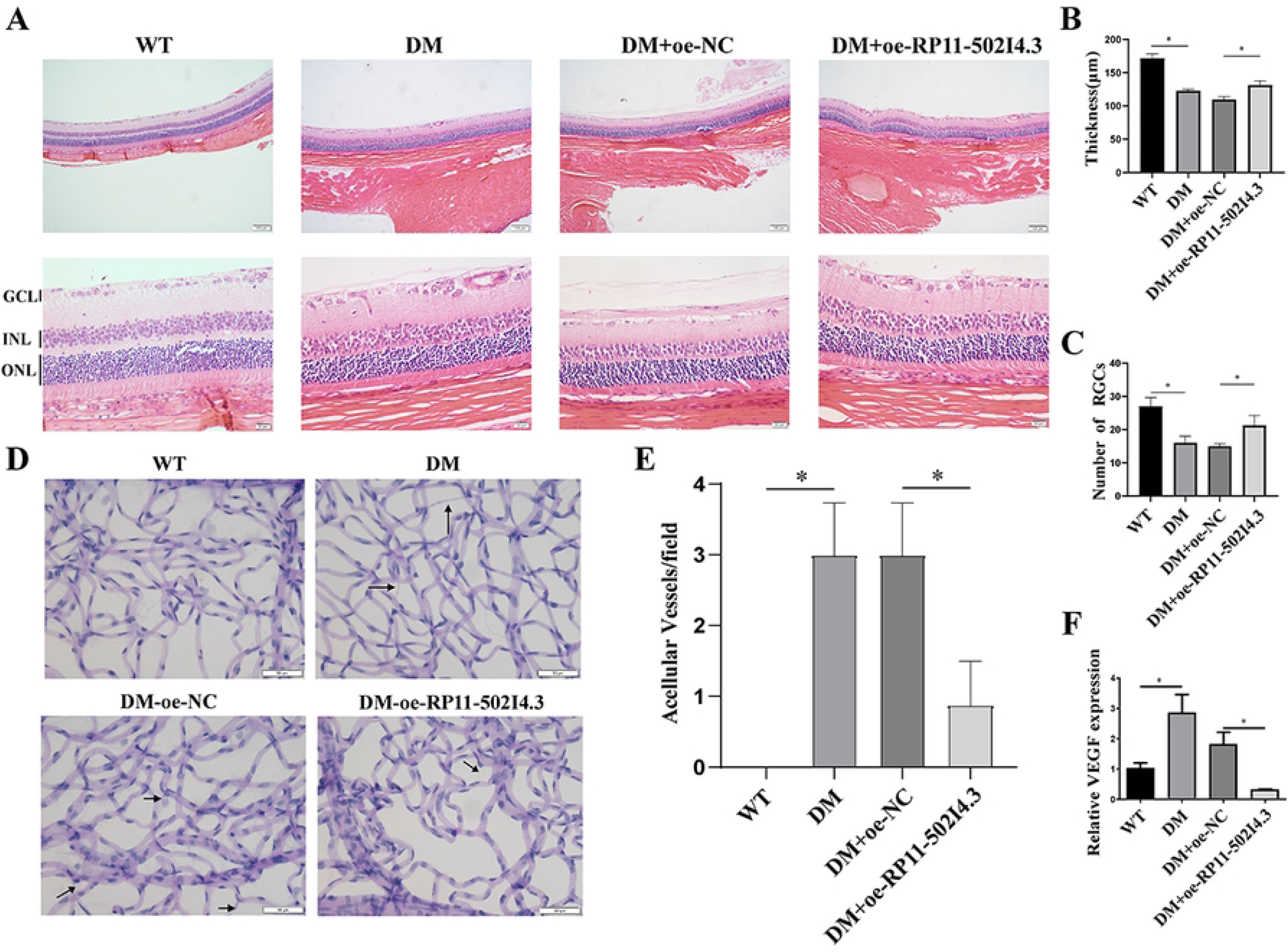

*In vitro* experiments demonstrated that the proliferation activity (Fig. 7A), migration count (Fig. 7B and C), and tube formation ability (Fig. 7D-F) of HG-induced HRMECs were increased compared with normal glucose conditions. However, these changes were delayed following RP11-502I4.3 overexpression. Similarly, VEGF expression was significantly downregulated in HG-induced HRMECs after RP11-502I4.3 overexpression (Fig. 7G).

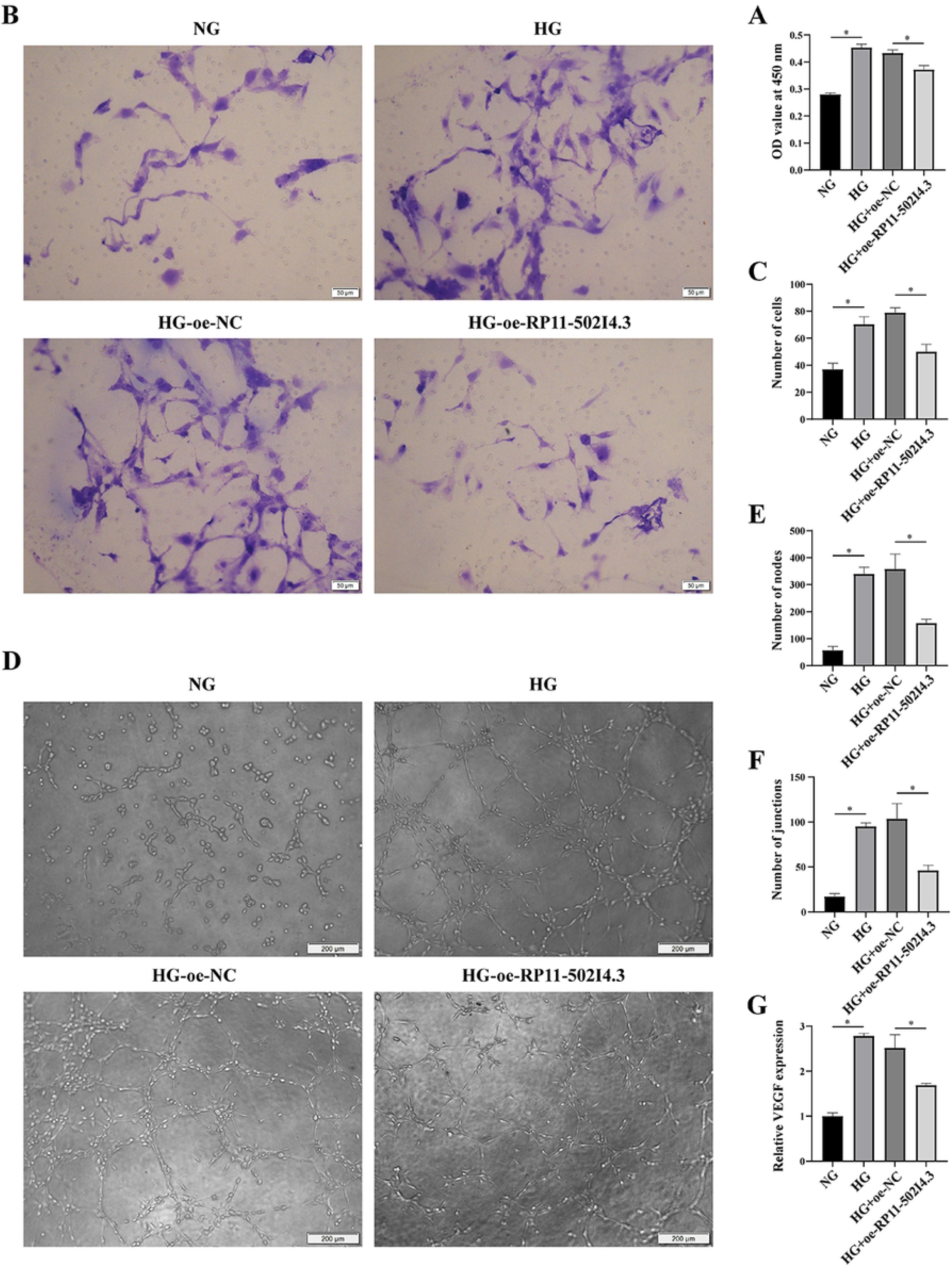

## Discussion

DR is the primary blinding eye disease in the elderly, with its pathogenesis being multifactorial[20]. A detailed understanding of this complexity is essential. Pericyte loss and EC apoptosis are triggered by chronic hyperglycemia, leading to retinal non-perfusion (RNP). As retinal hypoxia increases, VEGF expression rises, becoming a critical driver of PDR. This progression results in further RNP, which in turn enhances VEGF expression, creating a vicious cycle. Anti-VEGF therapy has become a vital component of DR treatment. Frequent anti-VEGF injections can disrupt this cycle, slow the progression of RNP, and improve retinopathy[21]. However, the frequency of injections in clinical practice is often low for various reasons, leading to disappointing visual acuity outcomes. Resistance to retinal neovascularisation remains a significant challenge. Gene therapy may be a new direction for the treatment of retinal neovascularization in the future.

LncRNAs have garnered significant attention in the study of vascular eye diseases and ocular angiogenesis[7]. Their functional mechanisms are diverse; some act as competitive endogenous RNAs, whereas others interact with proteins to regulate various angiogenic factors and numerous angiogenic and inflammatory signaling pathways[7]. LncRNAs are emerging as a novel way of treating DR[7, 20]. Recent studies suggest that lncRNAs offer additional layers of gene regulation owing to their diverse mechanisms[7]. In this study, RP11-502I4.3 was significantly downregulated in the hyperglycaemic HRMEC model, diabetic rat model, and VH samples from patients with PDR. RP11-502I4.3 may play a role in delaying retinal neovascularisation in DR; however, it remains poorly annotated and has yet to be reported in the context of DR or other diseases.

Many angiogenic factors are involved in retinal neovascularization. One of the most extensively studied of them is VEGF, which is a critical growth factor specific to endothelial cells (ECs)[7]. As a pathogenic mediator of retinal blood vessel leakage and abnormal growth, VEGF is a prominent therapeutic target in DR gene therapy research[22]. More and more studies have tried to intervene in the intraocular VEGF pathway. Dysregulated lncRNAs associated with models of ocular angiogenesis and can be categorized into pro-angiogenic lncRNAs (such as ANRIL, H19, HDAC4-AS1, SNHG16, HIF1A-AS2, HDAC4-AS1, Vax2os1 and Vax2os2) and anti-angiogenic lncRNAs (such as MEG3 and PKNY)[7]. In our study, RP11-502I4.3 overexpression downregulated VEGF expression. We hypothesize that RP11-502I4.3 downregulated may contribute to the formation of retinal neovascularisation by regulating VEGF.

DR represents a deterioration of the neurovascular unit (NVU), which comprises neurons, glial cells, and vascular cells. These components engage in intricate functional coupling and contribute to every stage of DR[23]. The molecular mechanisms underlying NVU dysfunction are multifactorial[24]. Numerous studies have investigated the factors associated with endothelial dysfunction. These investigations have primarily focused on the connection between ECs and retinal neovascularisation [25, 26, 27, 28, 29, 30, 31]. Angiogenesis is a complex multistep process[32]. Various reviews have indicated that ECs are the primary targets of hyperglycaemic damage[33, 34, 35, 36, 37]. Pericyte loss increases microvascular penetrability and exacerbates damage to the blood-retinal endothelial barrier. Long-term retinal microvascular damage leads to retinal ischemia[22]. Subsequently, VEGF upregulation promotes the proliferation, migration, and angiogenesis of ECs[38, 39]. Given the critical role of ECs in angiogenesis[38], we selected HRMECs as the target cells for this study. However, dysregulated lncRNAs can adversely affect various retinal cells[40, 41]. Except for ECs, other retinal cells can produce VEGF under HG conditions[24]. Almost all retinal cells can act as effectors or donors of VEGF and interact with each other through these mechanisms. The complex interactions among multiple factors highlight the intricate and complex nature of DR, indicating that treatment with a single factor is insufficient to reverse its progression[24]. Whether RP11-502I4 in HRMECs interacts with neighboring cells, such as pericytes, to induce endothelial dysfunction and contribute to the pathogenesis of DR remains unknown. Further research on other potential target cells is warranted.

Various *in vivo* models of DR have been developed for experimental research[42]. We selected rats as the animal model. However, they do not fully reappear all the characteristics of DR, such as microaneurysms and neovascularisation[43]. Only a few higher-order animal models can mimic the retinopathy observed in the late stages of DR[44]. Despite these limitations, rodent models provide valuable insights into the pathogenesis of DR[43].

Gene therapy is designed to deliver a therapeutic transgene. Luxturna, a virus-based gene therapy for Leber’s congenital amaurosis, was approved, marking a significant milestone in ocular gene therapy[45]. Gene therapy for polygenic eye diseases, including DR, has garnered increasing interest. Non-coding RNA delivered through viral vectors has shown a potential to inhibit the progression of DR effectively[2]. Gene therapy has several development avenues, such as gene augmentation, gene-specific targeting, and genome editing. Currently, gene therapy for DR primarily focuses on gene-specific targeted therapies, which require the construction of viral vectors to overexpress the target gene. Although viral vectors have been used in treatment, their use still needs further study[46]. New molecular-targeted drugs and drug delivery systems should receive more attention[46]. Emerging technologies for RNA therapy, such as nanoparticle-conjugated lncRNA overexpression and small interfering RNA-based lncRNA knockdown systems, suggest that lncRNA treatments may soon become a reality.

## Limitations

Our study had certain limitations. Firstly, because of the limited annotation of RP11-502I4, we were unable to predict its associated mRNA or signaling pathways using bioinformatics. Consequently, we focused on exploring the regulatory mechanisms of RP11-502I4 in retinal neovascularization. Secondly, only ECs were selected as target cells *in vitro* experiments. In the future, further investigation into the role of other retinal cells in this process is needed, which may help elucidate the potential connections and interactions between ECs and neighboring cells when RP11-502I4 inhibits retinal neovascularization. Finally, further research on the clinical application of lncRNAs is required to assess their curative effect.

## Conclusions

Therefore, we found that RP11-502I4.3 expression was lower in the retinal tissue of diabetic rats and HG-stimulated HRMECs compared with normal glucose conditions. The dysregulated RP11-502I4 is involved in the neovascularization of DR. This process may be mediated by VEGF.

## Supplementary Information

Not applicable.

## Acknowledgments

Not applicable.

## Author contributions

**Conceptualization:** Lan Zeng, Yuhao Wu.

**Data curation:** Lijuan Zhu.

**Formal analysis:** Junhao He.

**Funding acquisition:** Wei Tan.

**Investigation:** Lan Zeng, Yuhao Wu.

**Methodology:** Lan Zeng, Yuhao Wu.

**Project administration:** Wei Tan.

**Resources:** Wei Tan.

**Software:** Yuan Yuan.

**Supervision:** Lan Zeng, Yuhao Wu, Wei Tan.

**Validation:** Xiaocong Wang.

**Visualization:** Kai Tang.

**Writing – original draft:** Lan Zeng, Yuhao Wu.

**Writing – review & editing:** Wei Tan.

## Availability of data and materials

The datasets are available from the corresponding author upon reasonable request.

